# The effect of community dialogues and sensitization on patient reporting of adverse events in rural Uganda: uncontrolled before-after study

**DOI:** 10.1101/402503

**Authors:** Helen Byomire Ndagije, Leonard Manirakiza, Dan Kajungu, Edward Galiwango, Donna Kusemererwa, Sten Olsson, Anne Spinewine, Niko Speybroeck

## Abstract

**Background:** The patients that experience adverse events are in the best position to report them, only if they were empowered to do so. Systematic community engagement and support to patients in a rural setting to monitor any potential harm from medicines should provide evidence for patient safety.

**Methods:** This paper describes an uncontrolled before and after study aimed at assessing the effect of a community engagement strategy, the Community Dialogues and Sensitization (CDS) intervention between January and April 2017, on the knowledge, attitude and practice of reporting adverse drug events by community members in the two eastern Ugandan districts. A representative cross-sectional baseline household survey was done prior to the intervention in September 2016 (n=1034) and the end-line survey (n=827) in July 2017.

**Results:** After implementation of the CDS intervention, there was an overall 20% (95% CI=16- 25) increase in awareness about adverse drug events in the community. The young people (15- 24 years) demonstrated a 41% (95% CI =31-52) increase and the un-educated showed a 50% (95% CI=37-63) increase in awareness about adverse drug events. The attitudes towards reporting increased overall by 5% in response to whether there was a need to report ADEs (95% CI =3-7). An overall 115% (95% CI =137-217) increase in the population that had ever experienced ADEs was also reported.

**Conclusion:** Our evaluation shows that the CDS intervention increases knowledge, improves attitudes by catalyzing discussions among community members and health workers on health issues and monitoring safety of medicines.

## Background

Globally, adverse drug reactions account for up to 18% of hospital deaths 1-5. An adverse event is defined as any untoward medical occurrence in a patient administered a medicinal product and which does not necessarily have to have a causal relationship with the treatment, including worsening of the clinical condition6. Like many resource- limited countries, Uganda uses spontaneous reporting of suspected adverse drug events (ADE) predominantly by health workers to monitor the safety of medicines7. However, one of the biggest inherent limitations of this method is under-reporting. In as much as the Ugandan pharmacovigilance regulations of 2014 require healthcare professionals to report adverse drug events, the grossly under-resourced health sector makes it hard for health workers to report them8. A recent study reported an incidence of 25% of hospital-acquired suspected ADEs among Uganda inpatients9. Antibiotics and anti-malarials have been the most commonly implicated drugs in community-acquired ADEs in surveys of healthcare professionals10. With increased access and use of medicines in the community, it is becoming increasingly important to collect more safety information by involving the patients directly. However, healthcare professionals become less involved in patient treatment thus minimizing their role in healthcare delivery. The Ugandan National Pharmacovigilance Centre (NPC) has its main goal of promoting patient safety by monitoring adverse drug reactions. It intends to implement a program of a dialogue-based intervention aimed at encouraging community members to report and possibly contribute to prevention of adverse drug events and any drug-related issues in the community.

The Community Dialogue and Sensitization (CDS) approach was a hybrid modification of the community dialogue (CD) previously used to stimulate community support and engagement in the context of integrated community case management (iCCM) of childhood diarrhoea, pneumonia and malaria by Malaria Consortium in Zambia, Mozambique and Uganda^11^. The CDS model added a sensitization campaign using radio messages, posters and brochures to raise drug-safety awareness encourage dialogue and involve the community in designing solutions to pertinent issues. This approach involved a participatory communication process of sharing information through existing community-based structures and aimed to enable communities make informed choices and to take individual and collective action.

The CDS model assumed that the respondents knew the common diseases that affected the population across different ages and the treatment they often received. We assumed that the respondents knew the different common points of care in the community. Another assumption made was that the community members were familiar with the adverse events that they commonly experienced as well as the reporting channels in the community for the different service providers. This model assumed that respondents did not know that it was their right to report ADEs and their obligation to give feedback on ADEs to the national medicine regumatory authority to improve drug safety in the country. Some community members therefore had misconception about reporting ADEs and some patients accessed drugs through the most convenient avenues to them.

The messages developed for this study focused on informing communities that any drug was capable of causing ADEs and monitoring them was essential in improving drug safety. This paper describes an uncontrolled before and after study aimed at assessing the effect of a community engagement strategy, the Community Dialogues and Sensitization between January and April 2017, on the knowledge, attitude and practice of reporting ADEs by community members in the two eastern Ugandan districts

## Materials and methods

### Context

The CDS intervention was implemented in two predominantly rural districts of Iganga and Mayuge in Eastern Uganda. This area hosts a health and demographic surveillance site that works closely with the communities, health facilities and the district health office. The Iganga-Mayuge health and demographic surveillance site (IMHDSS) is run by Makerere University Centre for Health and Population Research (MUCHAP). The population here is a very young. About half is under 15 years and a total birth rate of five children. The IMHDSS population is largely homogeneous with 83% being from a single tribal-group, the Basoga. At the IMHDSS, there is bi-annual update of individual and household data conducted between February-May and August-November in the 65 villages. The updates contain information on pregnancy registrations and outcomes, in and out-migrations, births and deaths. Prior to each round, an information officer from each village gives feedback from the previous round and identifies the 35 data collectors for a 2-day training.

### The Intervention: Community Dialogue and Sensitization

The implementation team conducted a preliminary audience evaluation to understand the community needs and interests, gauge reactions to different communication strategies and identify preferences for various prototype materials. The National Pharmacovigilance Centre and MUCHAP jointly developed the CDS toolkit. The toolkit included a facilitators’guide- book with ten repeatable steps- key talking points, pictorial posters, and a monitoring and feedback tool. Tools and images were pre-tested with the target audience before finalizing the toolkit The CDS intervention included several components (See table 1) conducted between December 2016 and April 2017. The CDS meetings began after the official launch of the project. The goal of the community dialogue meetings was to improve the knowledge of the community members, about ADEs, improve attitude towards reporting ADEs and to minimize barriers to reporting ADEs. The primary target audience were the primary household member in charge of health care decisions, private health care providers, health workers at different levels, drug shop operators, pharmacies, district policy makers where possible, and patients.

**Table 1:**
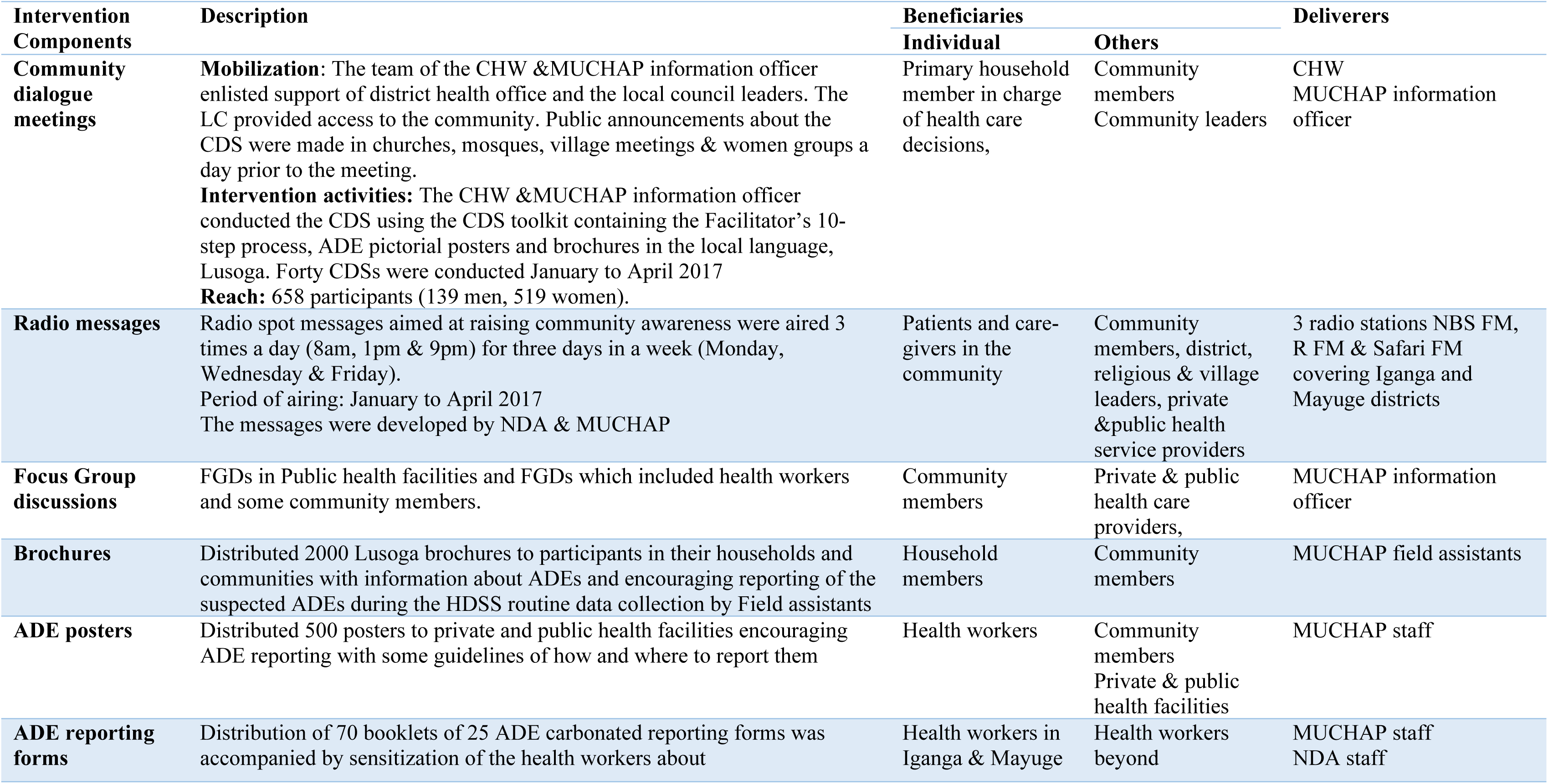
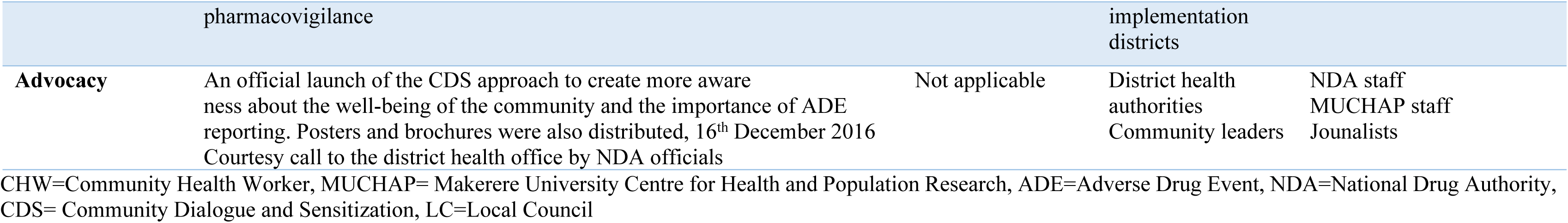
Multiple interventions used to improve patient reporting

After a courtesy visit to the district health office, the team explained the planned activities to the district health officer, district drug inspector and the district health educator. The teams went out in pairs composed of a community health worker and a note-taker from MUCHAP. The team enlisted the help of the local council heads to introduce them to the community to gain the acceptance of community members. Mobilization took the form of public announcements in churches, mosques, village meetings and women groups, a day before the CDS meeting. Announcement messages included the purpose, day, time and venue of the meeting. Teams emphasized to community members that the meeting would contain information that is important to their health and well-being.

During the community dialogue, a local leader or elder opened the session and introduced the team. The team leader then explained the purpose of the meeting and stimulated the dialogue into the ten-step process of the CDS toolkit and the talking points around ADEs. Community dialogues provided communities with the opportunity to discuss extensively within their villages, newly available healthcare services like patient ADE reporting. In addition, the members discussed how they could best benefit from the CDS approach and support it. In each village as far as possible, there were separate men and women’s meetings.

### Study design, respondents and sampling

This uncontrolled before and after study was conducted between September 2016 and August 2017 in the Iganga/Mayuge Health Demographic Surveillance Site (IDMSS). Prior to the CDS intervention in September 2016 and the end-line survey in July 2017, we conducted a representative cross-sectional-baseline household survey. The entire adult population of the study area was considered as the sampling domain, where all households were eligible for selection. The total sample size of 778 for each survey was calculated in order to allow for a comparison of proportions between two groups, i.e. baseline and end-line respondents. Assuming a baseline proportion with an acceptable level ρ of 0.5, and testing at the 0.05 level, a sample size of 389 for each survey was determined to give 80 per cent power to detect a change of at least ten per cent of the primary outcome. To allow adjustment for confounders, non-response and design effect, we doubled the sample size, to obtain the required sample size. Sampling involved a single-stage household sampling. In each of the 65 villages in the IMHDSS surveillance area, the study team sampled an equal number of households using a simple random sampling approach with the help of community leaders. For the purpose of the surveys, a household was defined as a group of people who routinely lived and ate together. One person per selected household was interviewed. The target for interviews was the person best placed to answer questions about the household’s health in the community members’ questionnaire (Additional file # 1). All community based health facilities in the intervention area were included in the survey. At least two healthcare workers from each health facility were randomly selected and interviewed using a specific healthcare provider questionnaire (Additional file # 2). The facilities considered were both private and public owned and these included hospitals, health centre VI-II, pharmacies, drug shops and shops that sell drugs with other merchandise.

### Data collection

A quantitative assessment of target communities’ knowledge, attitudes and practices of patient ADE reporting was done using identical questions before and after the CDS intervention.

Interviews were carried out by local field researchers, using a pre-tested structured and validated questionnaire translated in the local language (Lusoga). The field researchers were instructed to read out the survey questions exactly as rendered on the questionnaire. Instructions for field researchers with regard to whether the question required a single response or whether multiple responses were possible and answer options were provided in the questionnaire in English. Answer options were not read out aloud and field researchers were instructed not to suggest answers to the respondent. An ‘other’ category was provided for most questions and field researchers were instructed to note down respondents’ answers if they could not clearly assign the answer to an existing answer category.

All field team members had previous experience of conducting or supervising field research. Field researchers and supervisors attended a two-day training course covering data collection tools, field procedures and interview techniques. Field supervisors received an additional days’ training focusing on supervision of field teams as well as the sampling process. Following the pre-test, a half-day training session was conducted to discuss challenges identified during the pre-test. The training materials were prepared by MUCHAP and NDA and the training was conducted by MUCHAP, who were responsible for coordination and supervision during the field work. Field supervisors were tasked with monitoring the quality of the data collected and seeking clarification from the field researchers where necessary. At the end of each day, they were responsible for conducting a feedback meeting with their team, giving researchers the opportunity to discuss and resolve challenges and providing feedback and training to the team as appropriate. They reported to the study coordinator daily, summarizing progress made, challenges encountered and discussing field work to be completed on the following day.

### Data entry and analysis

Data entry was done using EpiData 3.1 (EpiData Association) software by ten trained data entry officers. All records were double entered to ensure accuracy. First and second entries were done by different data entry officers for each village. Where differences between first and second entry were detected, data were verified by checking the record against the paper questionnaire. If in doubt, data entry officers were instructed to log their query and discuss it with the study coordinator. The data was transferred to STATA Version 12 (StataCorp LP) for further consistency checks and preparation for analysis. All percentages reported are population average estimates which have been adjusted to take into account the clustering of the study design. Responses recorded under ‘other’ by field researchers were reviewed and either re-assigned to an existing answer category, assigned to a new answer category or left in the ‘other’ category. All comparisons were done at 5% level of significance and 95% confidence intervals for the mean difference were constructed to test the significance of the difference before and after the intervention.

### Outcome measures

The primary outcomes measured comprised of the percentage differences between the knowledge, attitudes and practice of reporting before and after the CDS intervention. Knowledge was measured by the responses from the question “Do drugs “cause” negative or side effects?” For the respondents who answered “yes”, to the above question, their attitude towards reporting adverse drug effects was measured by asking if they considered it necessary to report an adverse drug effect. To understand the respondents’ practices, we asked them if they would report any ADE if encountered. The secondary outcome of this study was to establish the best ways to engage the community and this was included as part of the questionnaire during the surveys.

## Results

### Description of Survey respondents

There were 1034 respondents that participated in the baseline survey (before implementation of the CDS model) and 827 participated in the end-line survey. These were house-hold adult members who consented to our interview. All the participants that were recruited in the study consented to participate. There was a big positive difference in knowledge, attitudes and practices of respondents regarding reporting ADE between the baseline and end-line surveys. This is true though the number in baseline were more than those in end-line survey. Table 2 summarizes survey respondents’ profile in terms of age, education, religion and occupation.

**Table 2:** Community member description by social demographic characteristics

|  | Baseline n (%) N=1034 | End line n (%) N=827 |
| --- | --- | --- |
| **Age group** |  |  |
| 15-24 | 163 (15.8) | 137(16.6) |
| 25-34 | 304 (29.4) | 224(27.1) |
| 35-44 | 243 (23.5) | 200(24.2) |
| 45-54 | 145 (14.0) | 138(16.7) |
| 55-64 | 105 (10.2) | 54(6.5) |
| 65+ | 74 (7.2) | 74(9.0) |
| **Education** |  |  |
| None | 147 (14.2) | 90(10.9) |
| Primary | 532 (51.5) | 422(51.0) |
| Secondary | 310 (29.9) | 265(32.0) |
| Tertiary | 23 (2.2) | 27(3.3) |
| University | 22 (2.1) | 23(2.8) |
| **Religion** |  |  |
| Anglican | 246 (23.8) | 251(30.4) |
| Roman catholic | 93 (9.0) | 82(9.9) |
| Pentecostal | 103 (10.0) | 103(12.5) |
| Muslim | 573 (55.4) | 380(46.0) |
| Other | 19 (1.8) | 10(1.2) |
| **Occupation** |  |  |
| Professional | 29 (2.8) | 42(5.1) |
| Self employed | 562 (54.4) | 203(24.6) |
| Student | 24 (2.3) | 16(1.9) |
| Peasant | 175 (16.9) | 342(41.4) |
| Domestic work | 241 (23.3) | 127(15.4) |
| Other | 3 (0.3) | 96(11.6) |

### Knowledge about adverse drug reactions

After implementation of community dialogues about adverse drug events and reporting, there was an overall 20% increase in knowledge about ADEs in the community. Disaggregating knowledge of ADEs by background characteristics, revealed an even distribution positive change but higher (41.1%, 95% CI =31-52) among young people (15-24years), those with no education (50%, 95% CI=37-63) as shown in table 3.

**Table 3:** Knowledge of ADEs by respondent demographic characteristics before and after the CDS intervention

| Do drugs “cause” negative or side effects? |  |  |  |  |  |  |  |  |  |  |  |  |
| --- | --- | --- | --- | --- | --- | --- | --- | --- | --- | --- | --- | --- |
|  | Yes |  |  |  | No |  |  |  | DK |  |  |  |
|  | Before | After | % age diff. | 95% CI | Before | After | % age diff. | 95% CI | Before | After | % age diff. | 95% CI |
| **Age category** |  |  |  |  |  |  |  |  |  |  |  |  |
| 15-24 | 86 (52.8) | 102 (74.5) | 41.1 | 0.31-0.52 | 63 (38.7) | 33(24.1) | -37.73 | -0.57--0.19 | 14(8.6) | 2(1.5) | -82.56 | -1.22--0.43 |
| 25-34 | 173 (56.9) | 151 (67.4) | 18.5 | 0.10-0.27 | 110(36.2) | 67(29.9) | -17.40 | -0.32--0.03 | 21(6.9) | 6(2.7) | -60.87 | -0.82--0.39 |
| 35-44 | 145 (59.7) | 137 (68.5) | 14.7 | 0.06-0.24 | 84(34.6) | 55(27.5) | -20.52 | -0.36--0.05 | 14(5.8) | 8(4) | -31.03 | -0.50--0.12 |
| 45-54 | 79 (54.5) | 87 (63.5) | 16.5 | 0.05-0.28 | 51(35.2) | 43(31.4) | -10.80 | -0.30-0.08 | 15(10.3) | 7(5.1) | -50.49 | -0.76--0.25 |
| 55-64 | 54 (51.4) | 37 (68.5) | 33.3 | 0.18-0.49 | 37(35.2) | 13(24.1) | -31.53 | -0.59--0.04 | 14(13.3) | 4(7.4) | -44.36 | -0.80--0.08 |
| 65+ | 45 (60.8) | 46 (62.2) | 2.3 | -0.13-0.18 | 26(35.1) | 25(33.8) | -3.70 | -0.30-0.22 | 3(4.1) | 3(4.1) | 0.00 | -0.32-0.32 |
| **Education** |  |  |  |  |  |  |  |  |  |  |  | 0.00-0.00 |
| None | 50 (34.0) | 46 (51.1) | 50.3 | 0.37-0.63 | 79(53.7) | 37(41.1) | -23.46 | -0.43--0.04 | 18 (12.2) | 7(7.8) | -36.07 | -0.63--0.09 |
| Primary | 294 (55.3) | 273 (64.8) | 17.2 | 0.11-0.23 | 191(35.9) | 132(31.4) | -12.53 | -0.23--0.02 | 47(8.8) | 16(3.8) | -56.82 | -0.72--0.42 |
| Secondary | 199 (64.2) | 193 (72.8) | 13.4 | 0.06-0.21 | 95(30.6) | 65(24.5) | -19.93 | -0.34--0.06 | 16(5.2) | 7(2.6) | -50.00 | -0.68--0.32 |
| Tertiary | 19 (82.6) | 25 (92.6) | 12.1 | -0.06-0.30 | 4(17.4) | 2(7.4) | -57.47 | -1.09--0.06 | 0(0) | 0(0) | 0.00 | 0.00-0.00 |
| University | 20 (90.9) | 23 (100) | 10.0 | -0.02-0.22 | 2(9.1) | 0(0) | -100.00 | -1.40--0.60 | 0(0) | 0(0) | 0.00 | 0.00-0.00 |
| **Religion** |  |  |  |  |  |  |  |  |  |  |  | 0.00-0.00 |
| Anglican | 140 (56.9) | 174 (69.3) | 21.8 | 0.13-0.3 | 91(37) | 69(27.5) | -25.68 | -0.40--0.11 | 15(6.1) | 8(3.2) | -47.54 | -0.66--0.29 |
| Roman Catholic | 59 (63.4) | 56 (68.3) | 7.7 | -0.06-0.22 | 29(31.2) | 23(28) | -10.26 | -0.35-0.15 | 5(5.4) | 3(3.7) | -31.48 | -0.62--0.01 |
| Pentecostal | 65 (63.1) | 77 (74.8) | 18.5 | 0.06-0.31 | 30(29.1) | 25(24.3) | -16.49 | -0.40-0.07 | 8(7.8) | 1(1.0) | -87.18 | -1.40--0.34 |
| Muslim | 308 (53.8) | 244 (64.2) | 19.3 | 0.13-0.26 | 217(37.9) | 118(31.1) | -17.94 | -0.28--0.07 | 48(8.4) | 18(4.7) | -44.05 | -0.58--0.30 |
| Other | 10 (52.6) | 9 (90) | 71.1 | 0.42-1.00 | 4(21.1) | 1(10) | -52.61 | -1.24-0.19 | 5(26.3) | 0(0) | -100.00 | 0.00-0.00 |
| **Overall** | **582 (56.3)** | **560 (67.8)** | **20.4** | **0.16-0.25** | **371(35.9)** | **236(28.57)** | **-0.20** | **-0.28--0.13** | **81(7.8)** | **30(3.63)** | **-0.53** | **-0.64--0.43** |

ADE=Adverse Drug Event, DK=Don’t Know, CI=Confidence Interval

### Attitudes about reporting adverse drug effects

In response to whether there it was necessary to report the adverse drug effects, there was an overall increase of 4.6% after the implementation of the CDS intervention. The difference in attitude after the intervention is presented in table 4 by social demographic characteristics of age, education and religion.

**Table 4:**
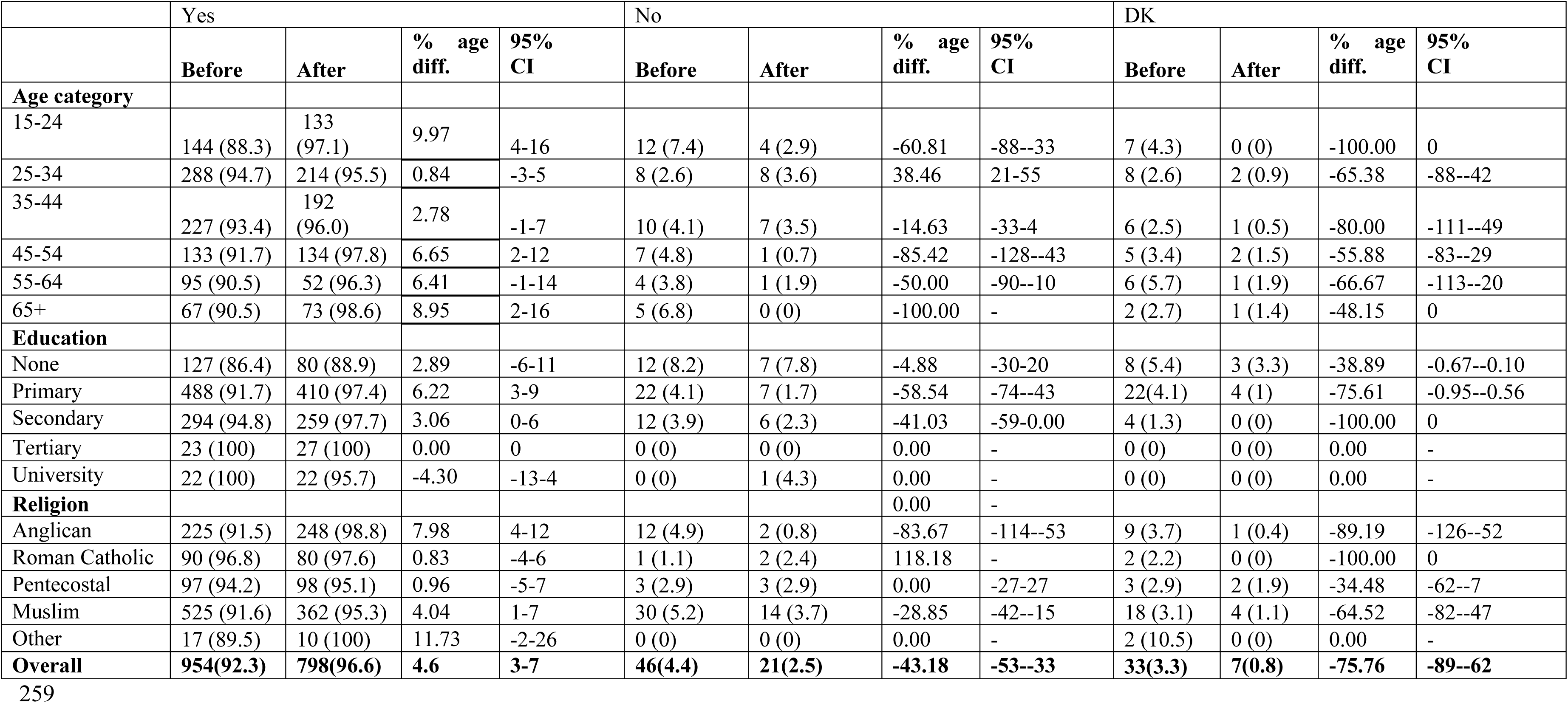
Necessity to report adverse drug events by social demographic characteristics before and after the CDS intervention

### Reporting ADEs by the respondents

While at baseline only 21% mentioned that they had experienced an ADE, this proportion more than doubled to 44% after the CDS intervention. However, there are variations in the different demographic groups, for instance, there was a reduction among those who had attained tertiary education level, as shown in table 5.

**Table 5:**
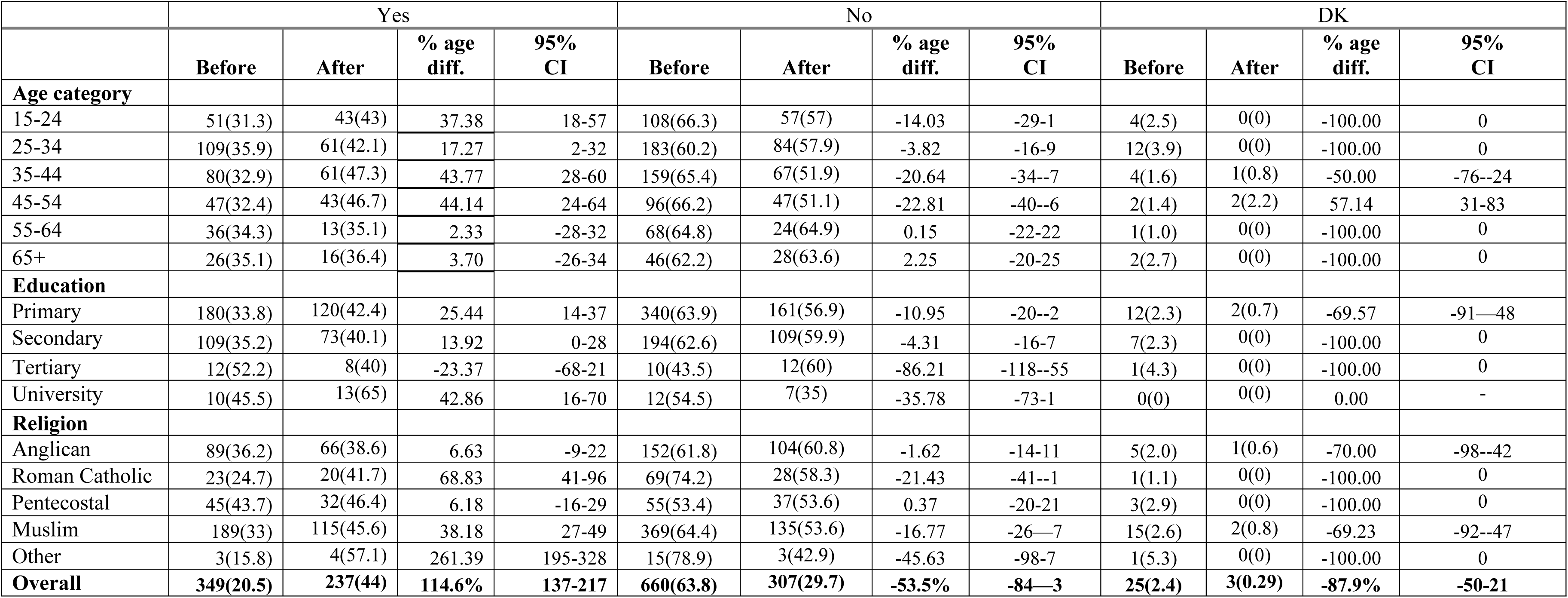
Reporting of ADEs by social demographic characteristics before and after the CDS intervention

|  | Yes |  |  |  | No |  |  |  | DK |  |  |  |
| --- | --- | --- | --- | --- | --- | --- | --- | --- | --- | --- | --- | --- |
|  | Before | After | % age diff. | 95% CI | Before | After | % age diff. | 95% CI | Before | After | % age diff. | 95% CI |
| **Age category** |  |  |  |  |  |  |  |  |  |  |  |  |
| 15-24 | 51(31.3) | 43(43) | 37.38 | 18-57 | 108(66.3) | 57(57) | -14.03 | -29-1 | 4(2.5) | 0(0) | -100.00 | 0 |
| 25-34 | 109(35.9) | 61(42.1) | 17.27 | 2-32 | 183(60.2) | 84(57.9) | -3.82 | -16-9 | 12(3.9) | 0(0) | -100.00 | 0 |
| 35-44 | 80(32.9) | 61(47.3) | 43.77 | 28-60 | 159(65.4) | 67(51.9) | -20.64 | -34--7 | 4(1.6) | 1(0.8) | -50.00 | -76--24 |
| 45-54 | 47(32.4) | 43(46.7) | 44.14 | 24-64 | 96(66.2) | 47(51.1) | -22.81 | -40--6 | 2(1.4) | 2(2.2) | 57.14 | 31-83 |
| 55-64 | 36(34.3) | 13(35.1) | 2.33 | -28-32 | 68(64.8) | 24(64.9) | 0.15 | -22-22 | 1(1.0) | 0(0) | -100.00 | 0 |
| 65+ | 26(35.1) | 16(36.4) | 3.70 | -26-34 | 46(62.2) | 28(63.6) | 2.25 | -20-25 | 2(2.7) | 0(0) | -100.00 | 0 |
| **Education** |  |  |  |  |  |  |  |  |  |  |  |  |
| Primary | 180(33.8) | 120(42.4) | 25.44 | 14-37 | 340(63.9) | 161(56.9) | -10.95 | -20--2 | 12(2.3) | 2(0.7) | -69.57 | -91—48 |
| Secondary | 109(35.2) | 73(40.1) | 13.92 | 0-28 | 194(62.6) | 109(59.9) | -4.31 | -16-7 | 7(2.3) | 0(0) | -100.00 | 0 |
| Tertiary | 12(52.2) | 8(40) | -23.37 | -68-21 | 10(43.5) | 12(60) | -86.21 | -118--55 | 1(4.3) | 0(0) | -100.00 | 0 |
| University | 10(45.5) | 13(65) | 42.86 | 16-70 | 12(54.5) | 7(35) | -35.78 | -73-1 | 0(0) | 0(0) | 0.00 | - |
| **Religion** |  |  |  |  |  |  |  |  |  |  |  |  |
| Anglican | 89(36.2) | 66(38.6) | 6.63 | -9-22 | 152(61.8) | 104(60.8) | -1.62 | -14-11 | 5(2.0) | 1(0.6) | -70.00 | -98--42 |
| Roman Catholic | 23(24.7) | 20(41.7) | 68.83 | 41-96 | 69(74.2) | 28(58.3) | -21.43 | -41--1 | 1(1.1) | 0(0) | -100.00 | 0 |
| Pentecostal | 45(43.7) | 32(46.4) | 6.18 | -16-29 | 55(53.4) | 37(53.6) | 0.37 | -20-21 | 3(2.9) | 0(0) | -100.00 | 0 |
| Muslim | 189(33) | 115(45.6) | 38.18 | 27-49 | 369(64.4) | 135(53.6) | -16.77 | -26—7 | 15(2.6) | 2(0.8) | -69.23 | -92--47 |
| Other | 3(15.8) | 4(57.1) | 261.39 | 195-328 | 15(78.9) | 3(42.9) | -45.63 | -98-7 | 1(5.3) | 0(0) | -100.00 | 0 |
| **Overall** | **349(20.5)** | **237(44)** | **114.6%** | **137-217** | **660(63.8)** | **307(29.7)** | **-53.5%** | **-84—3** | **25(2.4)** | **3(0.29)** | **-87.9%** | **-50-21** |

### Commonly reported adverse events

After the CDS intervention, there was an increase of more than 10% in the population who would consider reporting serious reactions (19%, 95% CI =16-21%), reactions to newly introduced drugs (15%, 95% CI = 11-18%), unexpected reactions (16%, 95% CI = 13-19%) and reactions due to herbal and conventional medicines taken together (20%, 95% CI= 16- 24%) as shown in table 6. Regarding the top most five ADE types that the respondents would report, it was found that serious reactions became more important after the CDS intervention. The rest of the order of importance for reactions to be reported largely remained unchanged from uncertain reactions to those of newly introduced drugs on the market, followed by unexpected reactions and reactions to drugs that have been on the market for long in descending order.

**Table 6:** A comparison of the type of adverse events that the respondents would report before and after the CDS intervention

|  | Yes/Agree n (%) |  |  |  | No/Disagree n (%) |  |  |  | Don't know n (%) |  |  |  |
| --- | --- | --- | --- | --- | --- | --- | --- | --- | --- | --- | --- | --- |
|  | Baseline | End-line | %change | 95% CI | Baseline | End-line | %change | 95% CI | Baseline | End-line | %change | 95% CI |
| Uncertain or suspected ADEs | 807 (78.0) | 721 (87.3) | 11.92 | 9-15 | 133 (12.9) | 84 (10.2) | -20.93 | -28--14 | 94 (9.1) | 21 (2.5) | -72.53 | -80--65 |
| Certain negative ADEs | 697 (67.4) | 634 (76.7) | 13.80 | 10-18 | 235 (22.7) | 168 (20.3) | -10.57 | -17--4 | 102 (9.9) | 24 (2.9) | -70.71 | -78--63 |
| Serious reactions | 797 (77.1) | 756 (91.5) | 18.68 | 16-21 | 143 (13.8) | 57 (6.9) | -50.00 | -57--43 | 94 (9.1) | 13 (1.6) | -82.42 | -90--75 |
| Mild reactions | 728 (70.4) | 573 (69.4) | -1.42 | -6-3 | 243 (23.5) | 246 (29.8) | 26.81 | 20-33 | 63 (6.1) | 7 (0.9) | -85.25 | -93--78 |
| Reactions to drugs which have been on market for a long period | 718 (69.4) | 649 (78.6) | 13.26 | 10-17 | 223 (21.6) | 146 (17.7) | -18.06 | -24--12 | 93 (9.0) | 31 (3.7) | -58.89 | -65--52 |
| Reactions to newly introduced drugs | 770 (74.5) | 706 (85.5) | 14.77 | 11-18 | 147 (14.2) | 73(8.8) | -38.03 | -46--31 | 117 (11.3) | 47 (5.7) | -49.56 | -57--42 |
| Common or well-known reactions | 639 (61.8) | 483 (58.5) | -5.34 | -10--1 | 285 (27.6) | 315 (38.1) | 38.04 | 32-44 | 110 (10.6) | 28 (3.4) | -67.92 | -75--61 |
| Unexpected reactions | 737 (71.3) | 683 (82.7) | 15.99 | 13-19 | 186 (18.0) | 107 (12.9) | -28.33 | -35--21 | 111 (10.7) | 36 (4.4) | -58.88 | -66--52 |
| Possible interaction with other drugs | 660 (63.8) | 582 (70.5) | 10.50 | 6-15 | 231 (22.3) | 179 (21.7) | -2.69 | -9-4 | 143 (13.8) | 65 (7.9) | -42.75 | -50--35 |
| Reaction due to herbal medicine | 573 (55.4) | 520 (62.9) | 13.54 | 9-18 | 370 (35.8) | 279 (33.8) | -5.59 | -11-0.00 | 91 (8.8) | 27 (3.3) | -62.50 | -70--55 |
| Reactions due to herbal & conventional medicine taken together | 643 (62.2) | 616 (74.6) | 19.94 | 16-24 | 301 (29.1) | 179 (21.7) | -25.43 | -31--19 | 90 (8.7) | 31 (3.7) | -57.47 | -64--51 |

ADEs = adverse drug events

### Best way to engage the community

The radio was reported as the best way to deliver the messages sensitizing the community members about ADE reporting among the respondents, whereas community meetings was regarded the best by health providers as shown in figure 1. The health-worker and the health facility was found to increasingly play a vital role of ADE sensitization among health workers and respondents. Residents of the community across the board were happy with the community dialogue meetings as a way of raising their awareness and as the best way to engage the community on ADE matters.

**Fig 1:**
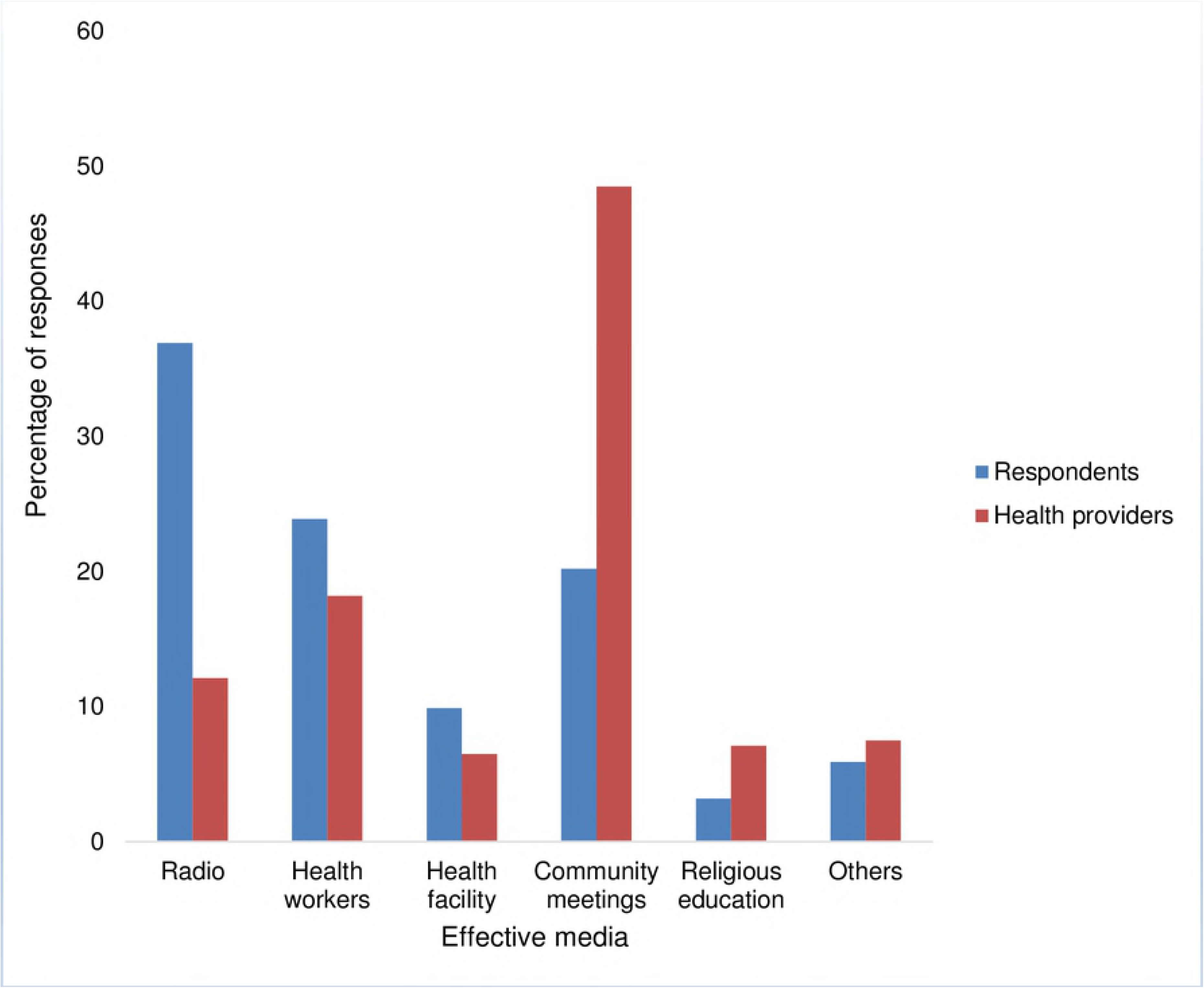
Comparison of options for reaching the community members with the ADE message between the community respondents and health providers.

## DISCUSSION

Our results suggest that the community dialogues and sensitization intervention increases knowledge, attitudes and reporting practices of adverse drug events across all demographic parameters in the rural communities. The population that reported having ever experienced an ADE more than doubled during the time of the CDS intervention. We also found that the intervention was widely acceptable through the focus group discussions and community meetings that we held. The local leaders were involved in mobilization and the local healthcare providers strengthened the messages on health consciousness and ADE awareness by the community. This study took advantage of the MUCHAP data collectors who were familiar with the community as they routinely collected data for the IMHDSS household surveys. The extent that such facilitated community meetings influence the quality of health care lies in the social capital that they raise. They provide an opportunity for networking and critical practical and emotional support, often leading to formulation of positive action plans and solidarity to action them^13^. A number of community dialogues and similar interventions have been implemented in similar settings but there is a paucity of data evaluating such interventions in the published literature. In line with results from our study, published evaluation data show positive effects of similar community-level interventions to improve awareness and attitude change regarding health issues^13^-^15^.

The results of this study indicated that, while the public is inclined to acquire information about ADEs and realize the benefits of reporting ADEs, their understanding of their essential role in reporting ADEs was insufficient. To increase awareness about ADEs, health workers reported that community meetings was the best method of raising their awareness and as the best way to engage the community on ADE matters while consumers suggested media campaigns on radios as the best way to deliver the messages sensitizing the community members about ADE reporting. One of the underlying assumption of the CDS intervention model was that individual exposure to the concept of monitoring and reporting ADEs would affect cognitions that continue to affect the behaviour of actual reporting of the events over a short term. Other studies have also reported the importance of media in raising community awareness and increase the likelihood of achieving new behaviour^15^, ^16^. This indicates that a combination of community dialogues and use of media campaigns on radios to sensitize people could increase reporting of ADEs in line with suggestions from other studies^15^-^18^. The present study demonstrated that the effect of CDS interventions would help improve ADE reporting. However, community interventions are time bound and hence continuous public educational programs on pharmacovigilance are essential to enhance reporting in the end. The long-term effect of especially the sensitization part of the model may operate through social and institutional pathways in addition to the individual learning that will require sustained levels of exposure through multiple channels over longer periods. Such effects tend to accumulate detectable change on certain sections of the audience over time and therefore should be assessed over time for a complete picture.

We found a significant improvement in attitudes of respondents towards reporting of ADEs. For example, in regard to whether it was necessary to report the adverse drug effects, there was an overall increase of 5% (95%CI =3-7%) after the implementation of the CDS intervention. There was an increase in the population who would consider reporting serious reactions, reactions to newly introduced drugs, unexpected reactions and reactions due to herbal and conventional medicines taken together. This showed that more than 15% of this community gained information about the negative effects of drugs in the short time of study. Similar results were found by other studies assessing educational interventions to improve attitudes of healthcare professionals and the adverse drug reaction-reporting rate in return^19^, ^20^.

Health providers and consumers’ willingness to report was reflected by an increase in the number of ADE reports submitted after the intervention. In regard to reporting practices, there was a change in respondents’ willingness to report serious reactions, reactions to newly introduced drugs, unexpected reactions and reactions due to herbal and conventional medicines taken together after the CDS intervention. The increase in willingness of health providers to report ADEs after community intervention was also reported by other studies^21^-^23^. However, the actual reporting could not be measured immediately since it’s an impact measure and therefore could not be ascertained immediately after the CDS intervention. The study results are still very informative for a policy on patient ADE reporting.

In Uganda, patients and consumers report adverse drug events either indirectly through their healthcare provider or directly through the newly established online reporting system^24^. In the rural setting, the capacity of patients to report was assessed by a proxy variable seeking to find out if the respondents had ever experienced or reported an ADE both before and after the CDS intervention. The 115% (95%CI =137-217) increase in recognition and reporting the ADE to the researchers is evidence that a separate system needs to be established for direct patient or consumer reporting. The launch of this system should be accompanied by a lot of sensitization delivered through multiple channels and with reporting tools that are tailored to the level of literacy and understanding of the rural community. There was no resistance to the intervention both at the community level and among the health workers. The increased reporting rate from this study and the wide acceptability of the CDS intervention in these two districts are ingredients for success of such a model of patient ADE reporting that can possibly be replicated in other limited resource settings. However, this successful implementation of the intervention in the study area could be because the respondents are used to research activities in the health demographic surveillance site and makes it hard to decline it. This means that, for a successful implementation of the same intervention in other settings, there is need for vigorous sensitization to prepare the communities and increase awareness.

Despite the initial increase of respondents’ and healthcare providers’ knowledge and attitudes towards ADE reporting following the CDS intervention, the influence of this intervention on the practice of ADEs reporting was not studied, which presents a major limitation in this study. The effect of the CDS intervention that was studied shortly after implementation, may not reflect the actual effect in the long term. Additionally, this study was conducted among healthcare providers and consumers from the rural settings from one region. Hence, the results of this study may be cautiously generalizable to rural settings in other regions.

The baseline and end-line surveys had limitations as to the CDS intervention model’s contribution to the changes in knowledge, perceptions and practice of patient ADE reporting given that randomly selected participants (before and after CDS) may, or may not have been beneficiaries of the project. There are also limits of the extent to which we can track direct changes to knowledge, practice and attitudes at an individual level given we did not track the same individuals throughout the life of the project.

There was no balance in samples selected at the baseline and at the end line surveys. This brought about overlap in frequencies and percentages by some demographic characteristics, for example in the gender and occupation variables. This could best be avoided by using exactly the same individuals before and after the intervention.

## CONCLUSION

In conclusion, the results of the current study showed that community dialogues and sensitization as a community intervention can increase knowledge, attitude and practices for reporting of ADEs, and that the respondents were able to apply the knowledge they gained from dialogues into their everyday life leading to increased reporting. Following the CDS intervention, the knowledge and attitude toward the ADE reporting seem to have improved. Hence, to improve ADE reporting among health-care professionals, there is a need to conduct periodic workshops and continued medical education frequently to sensitize them.

Despite having some fair knowledge that medicines have a potential to cause harm, the community shows signs of willingness to report the occurrence of adverse events through the preferred community channels, as well as to their healthcare providers. Some community members have experienced and reported ADEs to their healthcare providers. Further studies may be necessary to evaluate the impact of community interventions in the long-term effect after implementing the intervention. In its current state, the CDS intervention model is highly beneficial and can be adapted for consumer reporting in limited resource settings.

## DECLARATIONS

### Ethics approval and consent to participate

The study protocol was approved by the Mildmay Uganda research and ethics committee (REC REF 0604-2017) and Uganda National council of Science and Technology (HS 2247), the institution responsible for national research clearance in accordance with the World Medical Association Helsinki Declaration. Permission to conduct the study in the area was also obtained from NDA, the local administration of Iganga and Mayuge Districts, IMHDSS and MUCHAP. Written informed consent was obtained from the participants

### Consent for publication

Not applicable

### Availability of data and material

The datasets generated and analyzed during the current study are not publicly available but are available from the corresponding author on reasonable request.

### Author’s Contribution

HBN and DK designed the study. DAK made substantial contributions to conception, design of the study and acquisition of the data. LM, DK and HBN analysed and interpreted the data. LM was a major contributor in writing the manuscript. SO, AS and NS made substantial contributions during the review of the manuscript. All authors read and approved the final manuscript.

### Competing interests

Helen Byomire Ndagije, Leonard Manirakiza, Dan Kajungu, Edward Galiwango, Donna Asiimwe Kusemererwa, Sten Olsson, Anne Spinewine, Niko Speybroeck have no conflicts of interest that are directly relevant to the content of this study. This publication was supported by an agreement from National Drug Authority (NDA). Its contents are solely the responsibility of the authors and do not necessarily represent the official views of the National Drug Authority

### Funding

The study was supported with funds from the National Drug Authority.

## Acknowledgements

The authors express their profound gratitude to all patients and healthcare professionals who participated in this study. We are very grateful to the National Drug Authority for funding this study and to the Makerere University Centre for Health and Population Research team for letting us use the Iganga-Mayuge Health and Demographic Surveillance Site as a platform for data collection and data processing. Gratitude also goes to all research assistants who spent long hours during data collection and analysis. Finally, we thank the study participants in Iganga, Mayuge districts and the community leadership as well as the district authorities in those areas for allowing us to access all public health facilities at our convenience.

## Additional Files

Additional file # 1: Household survey questionnaire.doc Title of data:

Knowledge Attitudes and Practice (KAP) study on adverse drug reactions – Household questionnaire

Description of data

This was the questionnaire used to collect data about the households involved in the baseline and end-line surveys conducted before and after implementation of the community dialogues and sensitization.

Additional file # 2: Healthcare provider survey questionnaire.doc

Title of data:

Knowledge Attitudes and Practice (KAP) study on adverse drug reactions – Provider questionnaire

Description of data

This questionnaire was used to collect data about the healthcare providers in the community- based health facilities involved in the baseline and end-line surveys. At least two healthcare workers from each health facility were randomly-selected.

